# The FAIRSCAPE AI-readiness Framework for Biomedical Research

**DOI:** 10.1101/2024.12.23.629818

**Authors:** Sadnan Al Manir, Maxwell Adam Levinson, Justin Niestroy, Christopher Churas, Nathan C. Sheffield, Brynne Sullivan, Karen Fairchild, Monica Munoz-Torres, Sarah J. Ratcliffe, Jillian A. Parker, Trey Ideker, Timothy Clark

## Abstract

**Objective:** Biomedical datasets intended for use in AI applications require packaging with rich pre-model metadata to support model development that is explainable, ethical, epistemically grounded and FAIR (Findable, Accessible, Interoperable, Reusable).

**Methods:** We developed FAIRSCAPE, a digital commons environment, using agile methods, in close alignment with the team developing the AI-readiness criteria and with the Bridge2AI data production teams. Work was initially based on an existing provenance-aware framework for clinical machine learning. We incrementally added RO-Crate data+metadata packaging and exchange methods, client-side packaging support, provenance visualization, and support metadata mapped to the AI-readiness criteria, with automated AI-readiness evaluation. LinkML semantic enrichment and Croissant ML-ecosystem translations were also incorporated.

**Results:** The FAIRSCAPE framework generates, packages, evaluates, and manages critical pre-model AI-readiness and explainability information with descriptive metadata and deep provenance graphs for biomedical datasets. It provides ethical, schema, statistical, and semantic characterization of dataset releases, licensing and availability information, and an automated AI-readiness evaluation across all 28 AI-readiness criteria. We applied this framework to successive, large-scale releases of multimodal datasets, progressively increasing dataset AI-readiness to full compliance.

**Conclusion:** FAIRSCAPE enables AI-readiness in biomedical datasets using standard metadata components and has been used to establish this pattern across a major, multimodal NIH data generation program. It eliminates early-stage opacity apparent in many biomedical AI applications and provides a basis for establishing end-to-end AI explainability.

## 1. Introduction

Artificial intelligence (AI) readiness is a critical acceptance factor for biomedical datasets intended for use in ethical, FAIR (Findable, Accessible, Interoperable, Reusable) [1] AI applications, whether in the clinic or the laboratory. At the conclusion of an AI analysis, we must be able to explain and interpret the results [2–5]. This requires full transparency of the data preparation and analysis pipeline, including complete pre-model explainability of the input data and its various transformations, from patient or laboratory instrument to model training and execution [6]. Without substantiated explainability to establish epistemic validation for the foundational data, all further model building and analysis are subject to potentially catastrophic epistemic failure modes. Leonelli, who argues strongly against treating data as “ground truth”, stresses the role of understanding data provenance and characterization in determining its meaning, and hence its epistemic value in supporting subsequent interpretations [7].

FAIRSCAPE is a containerized client-server framework in Python, JavaScript, and React, with a human-in-the-loop AI assist mode. It was developed to provide transparency and validation with every biomedical dataset destined for Artificial Intelligence (AI) and Machine Learning (ML) analysis. It operationalizes AI-readiness criteria developed by the NIH Bridge to Artificial Intelligence (Bridge2AI) Standards Working Group [8] and integrates MLCommons Croissant [9], selected Croissant Responsible AI (RAI) [10], and LinkML schemas [11] for structured metadata representation. The framework continues to evolve in coordination with the Bridge2AI Standards Working Group, representing investigators from forty leading institutions, and through active collaboration with Global Alliance for Genomics and Health (GA4GH) [12], the NIH Generalist Repository Ecosystem Initiative (GREI) repositories consortium [13], the RO-Crate initiative at the University of Manchester [14], and the LinkML development team at Lawrence Berkeley National Laboratory to address emerging AI-readiness requirements across the biomedical AI community.

FAIRSCAPE creates deep metadata, including provenance graphs and dataset schemas, providing FAIR-compliant persistent identifiers (PIDs) for biomedical data and software. It creates and provides rich, human-readable Datasheets in HTML, extending the basic conceptions of Gebru et al. 2021 [15] to cover additional metadata required for biomedical AI-readiness. It also captures Model Cards [16] for cases where AI models are integrated with a data preparation pipeline, as in the Bridge2AI Functional Genomics Grand Challenge (Cell Maps for Artificial Intelligence - CM4AI) [17]. The framework supports a simple, direct upload process to any instance of the Harvard Dataverse, a repository within the NIH Generalist Repository Ecosystem Initiative (GREI). Interfaces to other GREI repositories are planned for the near-term.

Here we present the FAIRSCAPE client-server framework in detail. FAIRSCAPE establishes and validates pre-model AI explainability and AI-readiness of biomedical datasets by capturing rich information about the datasets and the software components involved in extracting and computing them, and managing this information in standardized exchange packages. The system has been validated by managing several large multimodal data releases in a major NIH program (Bridge2AI) in full conformance to that program’s AI-readiness criteria.

## 2. Related Work

### 2.1 FAIRness

The FAIR Principles established “Level 0” (Findable, Accessible, Interoperable, Reusable) requirements for reusable biomedical metadata and data. They are widely referenced and mandated but insufficiently specified for AI applications. FAIR Reusability subproperty R1.1 requires metadata describing a “plurality of accurate and relevant attributes” but does not specify which attributes are required. R1.2 states that metadata must be “associated with detailed provenance,” but does not specify which provenance model to employ and how detailed it must be. R1.3 states metadata must “meet domain-relevant community standards,” but does not specify what those standards are and places the burden of development on each domain, which is not appropriate for an overarching general standard.

Our approach operationalizes ethical, FAIR, and reusable metadata for the biomedical AI domain, guided by a well-defined set of criteria.

### 2.2 Digital Commons Environments and Generalist Repositories

Multiple digital commons environments have been developed over the past several decades. Specialist repositories, such as Sequence Read Archive (SRA), Database of Genotypes and Phenotypes (dbGaP), Gene Expression Omnibus (GEO), NHLBI BioData Catalyst, NHGRI Genomic Data Science Analysis, Visualization, and Informatics Lab-space (AnVIL), Genomic Data Commons (GDC), and others developed by NIH [18–21], and a multitude of resources developed at EMBL-EBI [22] are essential and widely used. They cannot, however, be used for datasets containing Protected Health Information (PHI).

One notable provenance-enabled data commons based on executable research objects was the NSF-funded Whole Tale project, developed by Chard et al. at the University of Chicago [23]. Whole Tale was designed to capture all data and software used in a research study for reproducibility. Unfortunately, this system does not appear to be actively maintained.

Robust repositories in the NIH GREI, including Dataverse, Zenodo, Figshare, OSF, Vivli, Dryad, and Mendeley [24–31], are widely used, including for multimodal data. The GREI consortium has published recommendations on adopting common metadata standards [32] based on the DataCite metadata schema [33]. We incorporate these key DataCite metadata elements into our framework and push FAIRSCAPE-packaged CM4AI datasets to the University of Virginia Dataverse instance for the long-term sustainability required by NIH. Unfortunately, at present, there is no mechanism to view our detailed metadata in Dataverse, other than viewing release-level information. Datasets and their schemas and provenance graphs must be exported from Dataverse for access to that information.

### 2.3 Provenance Metadata

Two notable and widely used specifications for provenance are the W3C PROV [34] and PAV (Provenance, Authoring, and Versioning) [35] ontologies. Developed by workflow experts to align outputs across heterogeneous pipelines, the W3C PROV ecosystem is grounded in twelve formally defined ontology standards of the World Wide Web Consortium [36–38].

PROV defines provenance as formal interactions among core Entities, Activities, and Agents, while allowing flexibility for domain practitioners to specify these components more deeply. PAV is intentionally lightweight and omits activities, resulting in a single-level provenance graph that specifies only “pav:derivedFrom” predicates on digital objects.

Seminal work leading to the EVI profile of PROV was the Micropublications ontology [39], which formalized claims and their support in biomedical articles as forms of bimodal defeasible argumentation. EVI reduced that scope to focus on datasets and their preceding computations, software, and input datasets, treating them as assertions justified by their provenance.

### 2.4 CEDAR Biomedical Metadata Templates

The Center for Expanded Data Annotation and Retrieval (CEDAR) metadata ecosystem [40,41] supports the creation of domain-specific metadata templates (reporting guidelines) for biomedical samples, protocols, and experimental entities, and is integrated with practically all relevant domain ontologies via the National Center for Biomedical Ontology’s BioPortal [42]. CEDAR has experienced widespread adoption for formalizing ontology-based characterization of biomedical experiments using domain-specific templates. CEDAR templates have been deployed to HuBMAP (Human BioMolecular Atlas Program) [43,44] to build over 30 metadata templates for use in that NIH program. Most notably, the NIH HEAL initiative [45] mandates its use to characterize experiments in over 1,000 projects. Overall, CEDAR has highly impressive adoption, with more than 5,500 registered users and 4,700 metadata templates created [46].

CEDAR’s strengths are complementary to FAIRSCAPE. While CEDAR does not connect its templates to deep provenance graphs in the data preparation pipeline and was not designed for AI-readiness preparation and evaluation against formal readiness criteria, it may offer an attractive complementary solution for templating rigorous reporting guidelines for ontology annotation of experimental materials and methods in the data preparation pipeline. Current practice in our program is to add ontology references of these entities ad hoc. This suggests a valuable role for CEDAR templates in a future integration project.

### 2.5 Other Domain-Specific Metadata Efforts

Other domain-specific metadata efforts in the life sciences include: the dmdScheme R package [47], ISA-Tab [48], and the Ecological Metadata Language (EML) [49]. The dmdScheme package was archived in 2023 and has not had the significant impact of CEDAR. ISA-TAB is used for metadata submission in several biomedical journals, but does not address AI-readiness, nor does it use a contemporary packaging standard such as RO-Crate. EML is a dataset-level documentation standard in the domain of ecosystem research that provides XML descriptions of datasets: who collected them, where, and by what methods. It forms the backbone of several important ecological repositories.

None of these three additional methods provides or evaluates AI-readiness, packages datasets in a portable exchange format, or provides resolvable deep provenance graphs.

### 2.6 Croissant and Croissant Responsible AI Metadata

The Croissant metadata vocabulary developed by the industry-sponsored MLCommons consortium provides essential metadata limited to computational AI-readiness. Croissant capabilities for computation-related metadata enable machine learning models to acquire **information** needed to train and be implemented on appropriately annotated datasets. However, Croissant views pre-model explainability merely as simple data engineering, as if the data provided were ground truths. This is demonstrably not the case, particularly in biomedical research due to its ethical requirements and often complex data extraction, transformation, analysis, and inference data preparation pipelines.

Although the Croissant RAI’s vocabulary introduces elements such as recommended and prohibited use cases to address this gap, it is a general-purpose specification that often lacks adaptation to the nuances of biomedical use cases. We therefore use it selectively. While some RAI properties can be repurposed for biomedical AI, a currently non-existent domain profile is required for several. The development of biomedical-specific extensions represents an important direction for future work. Most critically, RAI fails to deal comprehensively with data characterization and, critically, *treats data as presumptively given ground truths without detailed history*. As noted above, this can lead to substantial epistemic errors in making inferences, classifications, and predictions from data, particularly of the pervasive “Clever Hans” type [50– 53], with an impact on the scientific integrity of findings and ethical applicability in the clinic.

### 2.7 The Language Engineering Approach

A related but substantially different approach to documenting pre-model dataset characteristics is DescribeML [54], a domain-specific machine-readable language with associated tooling, provided as a plugin to Visual Studio Code. DescribeML structures textual metadata descriptions across dimensions of structure, provenance, and ethical/social risks to enable automated analysis. However, its provenance representation is relatively flat and non-resolvable, typically confined to statements such as “source: CIFAR10 from Kaggle” or “collectedBy X from Y”. While suitable for simply derived datasets, such as those used for online benchmarking challenges (e.g., Kaggle), this representation does not support biomedical applications or deep biomedical provenance and does not permit resolution to specific datasets, software, models, or responsible agents. The framework also includes metadata elements such as annotator demographics, which raise broader questions about consistency and scope in demographic reporting across roles in the research lifecycle. These elements were subsequently incorporated into Croissant RAI, which drew conceptually from DescribeML.

### 2.8 Machine Learning Specific Metadata Profiles and Checklists

As pointed out and analyzed in detail by Edmunds et al. 2026 [55], there is a growing, confusingly large body of work specifying standardized domain-specific ML metadata standards and checklists. These models, including the Data, Optimization, Model, and Evaluation (DOME) recommendations for ML in biology, focus primarily on AI/ML model design, training, and deployment rather than on data preparation pipelines or AI-readiness. While this body of work is essential for responsible model development, it does not address the foundational epistemic validation and data provenance that underpin those models – issues that FAIRSCAPE is designed to comprehensively support.

### 2.9 RO-Crate Packaging Tools

While tools like Describo [56] and its successor Crate-o [57] have successfully lowered the barrier to entry for building RO-Crates through general-purpose, user-friendly interfaces, they primarily rely on manual data entry and lack domain-specific validation.

FAIRSCAPE is not primarily about building generic RO-Crates. It is centrally concerned with producing ethical, FAIR, epistemically valid pre-model AI data. RO-Crates, in this context, are an affordance, not a *raison d’être*. FAIRSCAPE builds upon the RO-Crate standard but introduces an automated, enterprise-level architecture specifically tailored to generate and enforce the rigorous epistemic and ethical requirements of biomedical AI-readiness.

## 3. Methods

### 3.1 Development Approach

FAIRSCAPE was developed iteratively in multiple versions using agile methods over the past eight years, working closely with clinician-scientists, biostatisticians, laboratory investigators, ethics experts, and ontologists. We began by adapting a previously proven provenance-aware analytics framework for machine learning in the University of Virginia Neonatal Intensive Care Unit (NICU) [58,59] to support the distributed forty-institution research program Bridge2AI. The initial focus was on supporting CM4AI’s four separate contributing data production laboratories across three University of California campuses. This required client-side data and metadata packaging, for which we adopted the RO-Crate 1.2 specification while it was still in draft form, and initiated ongoing discussions with the RO-Crate development team. In Year 3 of the program, we began alignment with the AI-readiness criteria during their development.

#### 3.1.1 Iterative Standards Aligned Development

The AI-readiness criteria, which served as the basis for our metadata, were developed iteratively by experts in genomics, proteomics, clinical informatics, ontologies, machine learning, and metadata standards, across 17 institutions. We evolved our software in close alignment with the criteria and metadata mappings as they were developed. We also worked in close collaboration with data production groups in multiple institutions to validate the practicality of proposed metadata instantiations of the criteria. Software was iteratively validated across successive complex data releases in major NIH-funded projects.

Croissant and selected Croissant Responsible AI (RAI) mappings were added to facilitate interoperability with ML ecosystems. LinkML mappings were added to enable later semantic enhancement.

We found Croissant RAI’s “one-size-fits-all” metadata to be too general in many instances for adoption in biomedical research, requiring a domain profile, which we intend to develop. A leading example of over-generality would be the property rai:personalSensitiveInformation, which is intentionally broad to cover diverse sectors like computer vision, NLP, and social science, but lacks the granular precision required for legally mandated de-identification and security protection of subjects in clinical research.

Evaluation of metadata across the final seven dimensions and twenty-eight criteria of AI-readiness [8] was initially implemented as a self-evaluation. An automated evaluation based on the RO-Crate package metadata was later implemented, improving the scoring rigor and addressing the problems inherent in self-assessment [60].

### 3.2 RO-Crate Metadata Architecture

RO-Crates compliant with the version 1.2 specification [61] serve as the packaging format for FAIRSCAPE archival and distribution, supporting multiple relevant standards. This version was adopted while still in draft form to support multi-modal datasets in file trees with dataset references to external repositories. This is important to preserve deposition of specialist data modalities in domain-specific repositories where these exist, while retaining the integrative AI-readiness metadata and packaging.

RO-Crates are lightweight standardized packages (“crates”) consisting of a file tree of 1 to n levels with defined slots for metadata in specific serializations and vocabularies. They constitute cross-domain Boundary Objects, allowing sufficient plasticity to meet the local needs and constraints of the various parties implementing, exchanging, and using them while preserving robustness across the biomedical practice and communications ecosystem.

Metadata in RO-Crate’s required *ro-crate-metadata*.*json* file is serialized in JSON-LD [62] using schema.org [63]. FAIRSCAPE adds provenance characterization in W3C PROV [34] and the EVI domain profile of PROV [64,65] to these principal vocabularies. EVI domain profiles subclass the principal PROV classes and predicates for biomedical research. For example, *prov:Entity* is profiled by *evi:Dataset, Software, Model, Reagent, Sample*, and *Instrument* subclasses; *prov:Activity* as evi:*Computation, Experiment*, and *Annotation;* and *prov:Agent* as *evi:Person, SoftwareAgent*, and *Organization*. All *Entity* classes are resolvable to archived digital components and may be annotated with domain ontology terms.

FAIRSCAPE’s RO-Crates are composed of a top-level Release Package containing standard *ro-crate-metadata*.*json* and .*html* files, and a human-readable Datasheet that substantially extends the Gebru Datasheets for Datasets concept [15]. Along with the Datasheets, an AI-readiness evaluation histogram evaluates the seven dimensions of AI-readiness [8] across the full depth of the RO-Crate, and a LinkML schema translation to enable later deep semantic characterization via ontology mapping. The human-readable Datasheet is a textual and graphical rendering of the *ro-crate-metadata*.*json* file.

Sub-crates (sub-directories) in the file tree are organized by data modality, e.g., imaging, clinical records, genomics, proteomics, etc. Within each sub-crate are 1 to n dataset folders, each containing 1 to n files, all with the same format and construction. Dataset structure is characterized using JSON Schema [66] with Frictionless Data [67] validation.

Each component includes additional associated descriptive metadata using the schema.org vocabulary, plus PROV/EVI and optional ontology terms. Users may extend the metadata they associate with RO-Crates. FAIRSCAPE-CLI uses the Frictionless Data framework to generate JSON Schema definitions for tabular and HDF5 files, which are associated with their referenced datasets. Validation utilizing Frictionless ensures that datasets conform to their provided schemas. Every RO-Crate component receives a locally unique key. Data may be packaged directly or simply referenced using a Uniform Resource Identifier (URI) [68]. Because RO-Crate packaging occurs client-side and does not require server mediation for metadata construction, FAIRSCAPE avoids tight coupling to a specific server instance, enabling portability, archival independence, and long-term sustainability of AI-ready datasets. Once an RO-Crate is packaged, it may be uploaded directly to the server, where the local keys become resolvable ARK persistent IDs [69,70].

FAIRSCAPE manages the instantiation, transfer, archival, persistent identification, permissioning, and search of the RO-Crates, which are pre-validated on the client side using a Pydantic schema [71] and Frictionless Data [67,72]. Alternatively, if desired, the standard RO-Crate validator may be used with a preprocessor.

**Figure 1.** illustrates an example RO-Crate from a functional genomics dataset.

**Figure 1:**
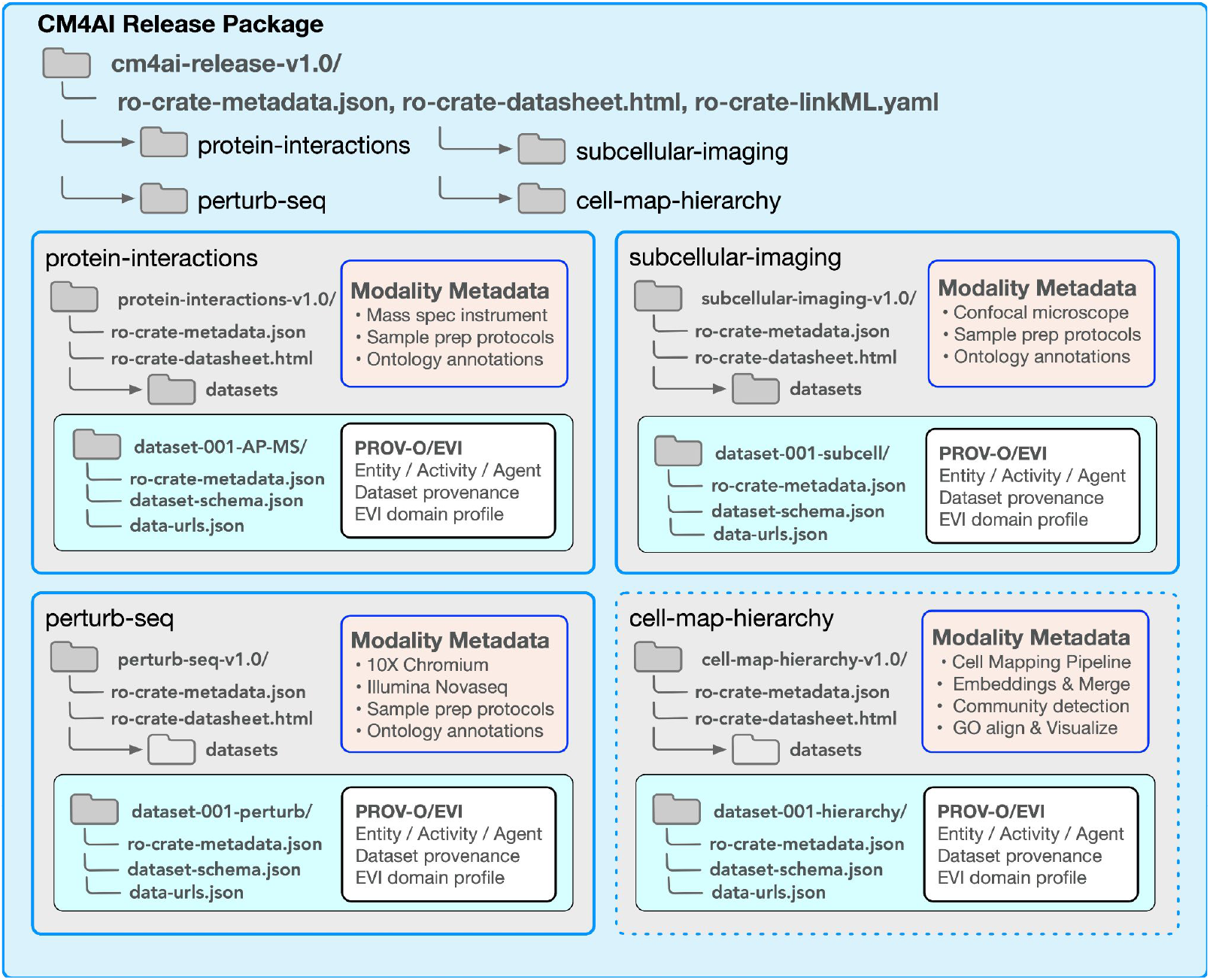
Cell Maps for Artificial Intelligence (CM4AI) RO-Crate schematic. All CM4AI release packages follow the RO-Crate 1.2 specification with some optional component additions. The top-level (root) crate is the “Release Crate” holding general information for the entire release. It includes a standard ro-crate-metadata.json file serialized in JSON-LD with schema.org vocabulary plus annotations from biomedical ontologies; an HTML rendered Datasheet; a LinkML translation of the Datasheet metadata; and links to sub-crates (shown in grey) for each modality in the release. CM4AI releases currently consist of 3 modalities (protein interactions, subcellular imaging, and perturbSeq), with the 4th modality (cell map hierarchy) planned for late 2026 shown in dotted outline. Each modality sub-crate contains 1-n dataset directories, shown in turquoise. These may contain 1-n files, a JSON Schema for the files, and the provenance graphs for the files.

## 4. Results

### 4.1 Features

The FAIRSCAPE framework generates, packages, and integrates descriptive metadata to robustly operationalize the AI-readiness criteria [8]. These metadata fully cover criteria across FAIRness, Provenance, Characterization, Pre-model Explainability, Ethics, Sustainability, and Computability. All metadata are represented in extended, human- and machine-readable Gebru-style Datasheets sealed with cryptographic signatures and packaged in lightweight standardized RO-Crate version 1.2 packages containing both the metadata and either (a) a link to associated datasets housed in an external sustainable repository, or (b) for smaller datasets, where allowed, the datasets themselves. RO-Crates are organized hierarchically as described above in Section 3.2, to support single-modality or multiple-modality data at shallow or deep levels of organization. The datasets are prepared client-side using either of three client packages, a command-line interface, or a simpler GUI application. Release-level metadata describing the entire package may be generated manually or via a human-in-the-loop AI assist.

The FAIRSCAPE framework comprises a set of Python and JavaScript packages, including alternative client versions (a command-line (CLI) Python 3 version, a GUI electron app, and a React-based JavaScript version) and a pip-installable, cloud-ready server that prepares datasets for AI analytics. AI-ready data packages present human- and machine-readable metadata characterizing the dataset(s) and link to, or include, those datasets. The electron GUI app is the simplest, intended mainly for less complex use with straightforward REDCap exports and analyses. The React-based JavaScript GUI provides all the necessary support for complex multimodal datasets, including human-in-the-loop AI assistance. The CLI client is designed for embedding in scripted pipeline code or in direct python calls by software engineers.

Data may be uploaded in simplified RO-Crate packages with minimal metadata, in compliance with DataCite and the GREI consortium recommendations. An automated evaluator notes deficiencies for various levels of requirement from “Basic” (DataCite schema only) to full AI-readiness, including Croissant metadata and LinkML translation. Datasets are automatically recognized by Frictionless Data libraries, and basic schemas are assigned with the option to expand them with user-defined data element names and descriptions. Alternatively, schemas may be imported from REDCap Data Dictionaries or specified in an attached JSON Schema description. FAIRSCAPE users specify deep provenance graphs on the client side using objects and predicates from W3C PROV and PROV’s EVI domain profile. The framework performs feature validation on uploaded data, software, and computations for all datasets.

FAIRSCAPE translates ethical, regulatory, schema, statistical, and semantic characterization of dataset releases, licensing, and availability information into formal human- and machine-readable metadata. It provides an automated AI-readiness evaluation, displayed as a histogram of the twenty-eight AI-readiness criteria, organized into seven axes.

### 4.2 Software Implementation

The FAIRSCAPE framework architecture is shown in **Figure 2**.

**Figure 2.**
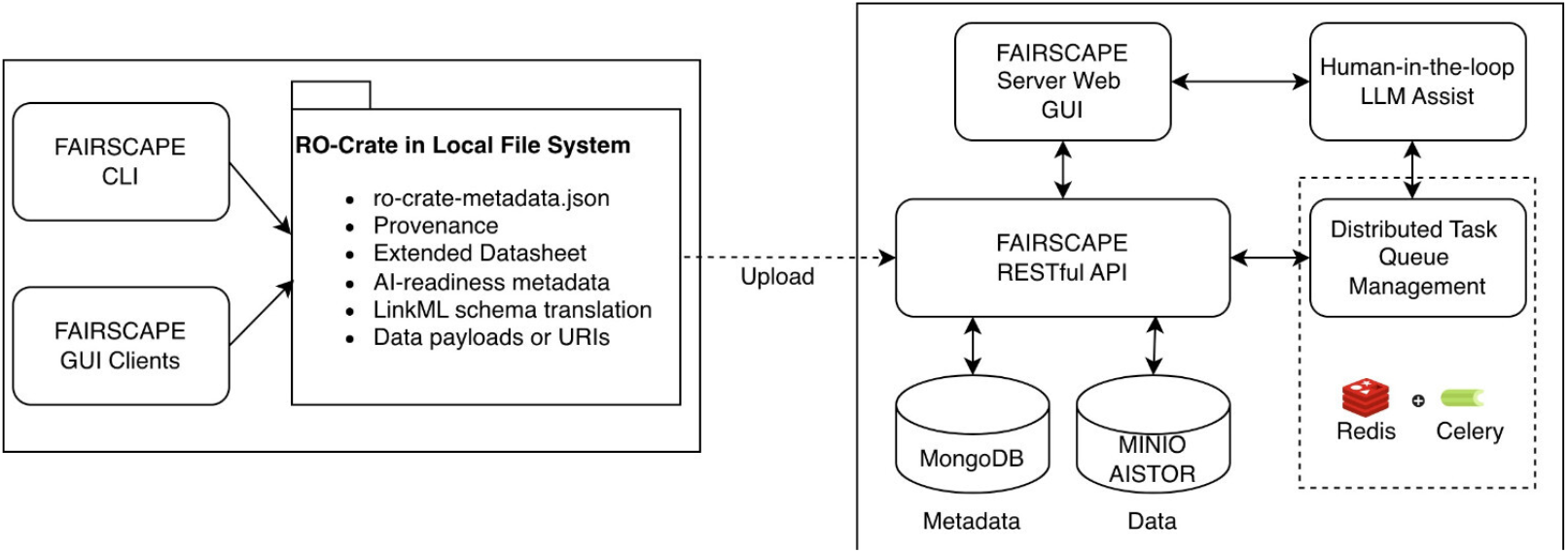
FAIRSCAPE framework architecture. The client-side applications package (meta)data in the canonical exchange layer as RO-Crates and submit to the server as compressed packages. The server REST API manages user sign-on with valid credentials, accepts the compressed packages, and passes management operations to the Redis message broker, which pushes metadata to the MongoDB store and datasets to the MinIO AIStor. Access to the metadata and data is controlled by user, group, project, and organization-level permissions stored in MongoDB.

#### 4.2.1 RO-Crate Packaging

RO-Crate packaging utilizes a combination of existing standards, as described in Section 3.2. The packaging hierarchy is flexible, except that the top-level or root RO-Crate must contain general “release” information describing the release and crate hierarchy, as well as a human- and machine-readable datasheet and a LinkML translation. We recommend including first-level sub-crates within the release crate, organized by data modality, and within each modality, by “dataset”, where a dataset is a folder of files with the same schema.

Packages may be prepared for a single computation or for a long series as required. It is not necessary to package each step for trivial or unimportant operations, as long as the complete software script performing the series is referenced and resolvable for inspection. In this way, FAIRSCAPE and EVI operate differently from a traditional workflow engine. RO-Crate-packaged computations may be connected in series, with Crate A as input to Crate B’s computations - the framework will then determine Crate A’s output dataset and connect it as input to the computation in Crate B to create a merged package.

#### 4.2.2 Client Software

Users may select the most appropriate client package for their use case: the Electron app is the simplest, the React GUI app is more complex and full-featured, and the command-line (CLI) version is more complex but supports in-pipeline package construction via calls from the command line or direct invocation as Python functions. Together, these tools create, manage, and upload RO-Crate packages to the server with detailed provenance graphs based on W3C PROV and the EVI Evidence Graph Ontology’s domain profile of PROV. FAIRSCAPE supports extending the metadata package, which is serialized in JSON-LD.

##### 4.2.2.1 FAIRSCAPE Command Line Interface Client

The FAIRSCAPE-CLI client is pip-installable and may be called either from the command line or directly as Python functions. It builds an RO-Crate and incrementally adds its components in a set of JSON-LD graphs using the following pattern:

- Computation <uses> Software
- Computation <uses> Model
- Dataset(1) <usedBy> Computation <generates> Dataset(2)

The underlying data model follows a defined provenance pattern in which a *Computation* entity *uses* specific *Software* (and optionally, *Model*) components, while a *Dataset* serves as input to the *Computation*, which subsequently *generates* an output *Dataset*. The tool allows users to define, infer, validate, and register schemas for common tabular data formats. Features for importing external datasets, such as NCBI BioProjects and Portable Encapsulated Projects (PEPs) [73] into RO-Crate formats, are also implemented. Additionally, it generates detailed evidence graphs that represent the provenance relationships among components, along with HTML datasheets for datasets in the release-level exchange package (top-level RO-Crate). Finally, the client provides mechanisms for RO-Crate release management by linking related sub-crates to support the packaging of multi-modal datasets and by facilitating publication to external repositories, including FAIRSCAPE and any instance of Harvard’s Dataverse.

##### 4.2.2.2 Electron Client

The FAIRSCAPE GUI client, built with Electron and JavaScript, guides the user through RO-Crate initialization and component upload. At each step, clients display a form to collect the required metadata, and the resulting JSON-LD is displayed on the side of the application. After completing all the required forms, users can review their created RO-Crate and its contents, package it into a zip file, and upload it to a FAIRSCAPE instance.

##### 4.2.2.3 JavaScript / React Client

An improved GUI client in React and JavaScript is also available, supporting both client human-in-the-loop AI-assisted and fully manual metadata packaging. The AI-assist feature instantiates significant release-level metadata in approximate form by providing documentation, other dataset metadata, publications describing data and methods, and a designed prompt, to a selected LLM. All metadata produced by the LLM are fully editable and require human approval with digital signoff before final acceptance.

#### 4.2.3 FAIRSCAPE Server

The FAIRSCAPE server is a cloud-ready Python application for managing metadata and data uploads in compressed format, with user-, group-, and project-based permissioning. It supports and manages coordinated ingress, access to, and publication of datasets via GUI and/or REST API. Enterprise-level FAIRSCAPE runs in a Kubernetes instance across multiple pods in our research IT environment, but it is flexible in its deployment targets and can be run on much smaller systems, including single machine deployments.

The FAIRSCAPE server receives, catalogs, indexes, and stores uploaded RO-Crate zip packages, extracts and registers their components and associated metadata, and stores that information. It uses the FastAPI framework and provides REST API access.

Storage is coordinated between a Mongo NoSQL database for metadata and a MinIO AISTOR object store for datasets. A worker task intercepts and schedules all requests to these data stores using a Redis cache as an in-memory message broker, allowing non-blocking submission of very large datasets.

All packaged datasets and external datasets or software referenced in another repository are managed in the MinIO AISTOR S3-compliant database.

The FAIRSCAPE Server provides a Web GUI for viewing metadata and downloading RO-Crates uploaded to FAIRSCAPE. After logging in, a dashboard displays all of the researcher’s uploaded RO-Crates, with links to download them or visit their landing page. Each landing page presents multiple views of an RO-Crate:

- A table summary of its required JSON-LD metadata and the files it contains.
- Serializations of its metadata in JSON-LD, RDF, and Turtle.
- An interactive visualization of dataset evidence graphs, built upon the React-Flow library, for exploring dataset provenance.

For cloud installations, MinIO access may be replaced by another S3 API. Metadata is managed in the Mongo NoSQL database. The server uses Redis as an in-memory cache and message broker to pass information and commands from the API to the internal Worker process for execution. Multi-user and group permissioning is handled as metadata. Objects stored in FAIRSCAPE may be pushed directly to any instance of the Dataverse academic repository system.

## 5. Discussion

As noted above, FAIRSCAPE was developed in close collaboration with laboratory researchers, clinician scientists, computer scientists, and standards experts across multiple programs using an agile method. Its design is highly responsive to the unique needs of multimodal biomedical datasets. The metadata elements of the framework are expected to evolve as additional needs are encountered.

One area of active development is adaptation to Croissant RAI. Many elements of this general-purpose standard, while widely promoted by the MLCommons consortium, are too general for biomedical research and thus require a domain-specific profile and further evaluation for proper use. For instance, current Croissant RAI properties suffer from implementation ambiguity, and *rai:annotatorDemographics* embeds ethical risks, such as promoting demographic essentialism (demography as determinant of scientific merit) and reinforcing power imbalances by over-emphasizing contributor identities. To address these shortcomings, a forthcoming biomedical domain profile will introduce formal subclassing, property selection, and alignment with domain-specific controlled vocabularies to ensure technical and ethical rigor.

An additional limitation and area for development concerns the AI-assisted evaluation of metadata. A number of release-level metadata elements are textual and therefore require qualitative human assessment to determine how well they align with the data they describe. Because such human assessments may introduce variability, we address this challenge in two stages. First, metadata components at the dataset release level undergo formal human review and digital signing by an authorized, PI-delegated curator (or the PIs themselves in smaller projects). Second, all criteria-mapped metadata are automatically evaluated using presence/absence and completeness metrics. Future work will incorporate AI-based assistants to suggest topic-specific improvements to descriptive metadata.

More broadly, AI explainability (XAI) is among the most rapidly expanding areas in the AI literature, yet it remains largely focused on post hoc explanations of trained models, with increasing but still limited attention to in-model approaches. This emphasis leaves a substantial epistemic gap, or “black box,” in the pre-model data preparation phase, which, in biomedical contexts, can involve complex and heterogeneous processes including variation in reagents, samples, instruments, lab software, and analytic pipelines. The growing integration of AI components directly into these upstream pipelines further amplifies the need for transparent, provenance-aware documentation prior to model training and deployment.

FAIRSCAPE closes this gap by providing substantial, verifiable pre-model explainability for biomedical AI projects, thereby laying the foundation for end-to-end explainability. In this sense, it extends Shapin’s concept of “virtual witnessing” [74,75], the practice established during the scientific revolution of publishing detailed methodological accounts to enable independent scrutiny, in contrast to the pre-scientific scholastic model of validation by simply citing authorities [76,77]. The rise of large-scale, automated data pipelines has complicated this norm by rendering many preparatory steps effectively invisible. FAIRSCAPE reconstitutes virtual witnessing in machine-readable form through detailed, resolvable documentation of data preparation processes. FAIRSCAPE’s RO-Crate packaging provides a robust Boundary Object [78–83] that mediates between domain practice groups in biomedical research, computer science, ethics, and ontology development. The deep, resolvable provenance graphs it provides allow data preparation pipelines to be replicated outside their initial context.

An additional development we are currently exploring is the provision of audience- or persona-focused textual explanations derived from FAIRSCAPE’s provenance graphs and related information in the exchange package.

Finally, GREI repositories are just beginning to consider AI-readiness issues. Integration of FAIRSCAPE tools, packaging, and viewers to support GREI AI-readiness for non-PHI information would be an extremely promising area for future development.

## 6. Conclusion

The FAIRSCAPE AI-readiness framework provides a rigorous, scalable, common platform for creating, managing, and distributing ethical, FAIR, AI-ready biomedical datasets. It was initially developed to provide deep provenance on computations in clinical predictive analytics applications in the University of Virginia Neonatal ICU. Subsequently, it was re-architected and significantly extended to support complete AI-readiness metadata packaging for diverse biomedical datasets, and validated across multimodal laboratory and clinical datasets in a major NIH-funded program, Bridge2AI.

FAIRSCAPE’s overall significance to biomedical research is in closing the epistemic gap in the pre-model data preparation phase — typically seen by purely CS-aligned AI researchers as limited to rote data engineering for model consumption. Taking biomedical data “as provided” as ground truth is not appropriate for biomedical research. The Bridge2AI program provided our team with an exemplary laboratory to develop methods to close this gap. FAIRSCAPE now provides foundational pre-model explainability metadata for this important purpose.

FAIRSCAPE remains under active development with the goal of broad applicability and adoption. The framework supports the production of FAIR, ethical, and epistemically robust AI-ready data. It supports all 28 practice-based AI-readiness criteria [8] across diverse biomedical contexts, with the potential to significantly strengthen the reliability and reusability of pre-model data for AI, and to increase the translational impact of AI applications across many research and clinical settings.

## CRediT Author Statement

SAM: Software, Supervision, Validation, Writing - Original Draft, Writing - Review and Editing; MAL: Software, Validation, Writing - Review and Editing; JN: Software, Validation, Writing - Review and Editing; CC: Software, Validation, Writing - Review and Editing; NCS: Software, Validation, Writing - Review and Editing; BS: Writing - Review and Editing; KF: Writing - Review and Editing; MMT: Funding acquisition, Methodology, Software, Validation, Writing - Review and Editing; SJR: Methodology, Writing - Review and Editing; JAP: Project Administration, Funding Acquisition, Methodology, Writing - Review and Editing; TI: Funding Acquisition, Methodology, Project Administration, Writing - Review and Editing; TC: Conceptualization, Funding Acquisition, Methodology, Project Administration, Supervision, Writing - Original Draft, Writing - Review and Editing.

## Declaration of Competing Interest

NCS is a consultant for InVitro Cell Research, LLC. Other authors declare no competing interests.

### Acknowledgements

This work was funded by the U.S. National Institutes of Health Bridge2AI program (OT2OD032742, OT2OD032701, 5U54HG012513-04), by the Eunice Kennedy Shriver National Institute of Child Health and Human Development (5R01HD072071-10), and by the University of Virginia Frederick Thomas Fund. The funders played no role in study design, data collection, analysis and interpretation of data, or the writing of this manuscript.

We thank the RO-Crate engineering team at the University of Manchester for very helpful discussions on RO-Crate architecture; the Bridge2AI Standards Working Group for collaboration on the metadata architecture; and Chris Mungall and staff at the Lawrence Berkeley National Laboratories Biosystems Data Science Department for developing the LinkML model and collaborating on a version of human-in-the-loop AI assist.

We are grateful to Katy Krahn for organizing the many effective meetings and discussions of the Center for Advanced Medical Analytics at UVA, which led to the initiation of this platform.

And we especially thank Carol Goble, Maryann Martone and Randall Moorman for many helpful discussions and interactions during platform design and development, and Mark Musen for discussions about potential CEDAR integration.

## Ethical Approval

The study conducted in University of Virginia Medical Center’s Neonatal Intensive Care Unit (NICU) used chart-reviewed Electronic Health Record (EHR) data. It was conducted under Institutional Review Board (IRB) Exemption 4 (45 CFR 46.104) because it involved secondary use of previously collected clinical data. All data in that study was anonymized.

## Declaration of Generative AI and AI-assisted technologies in the manuscript preparation process

During the preparation of this work the author(s) used Gemini 1.5 Pro in order to brainstorm potential image layouts, as well as to review the manuscript for compliance with journal editorial policies and to suggest potential improvements. The authors used Claude Opus 4.6 to point out grammar and capitalization issues in the pre-final copy. After using these tools/services, the author(s) created the original images, reviewed and edited the content as needed, and take full responsibility for the content of the published article.

## Data and Software Availability

Source code and Pydantic models for the FAIRSCAPE framework in this study are open-source, provided under Apache 2.0 license, and freely available on Github via Zenodo at the DOIs indicated below.

The LinkML code and documentation is freely available under Apache 2.0 license at the DOI indicated below.

Patient privacy constraints and Institutional Review Board (IRB) restrictions prevent the raw Electronic Health Record (EHR) data used for the NICU prediction task from being made publicly available.

A de-identified public dataset and an evidence graph for the NICU analyses are available on the University of Virginia Dataverse instance at the indicated DOIs below.

### Software and Tooling

- **FAIRSCAPE AI-readiness Framework:**
  - Levinson, M.A., et al. 2026 - FAIRSCAPE CLI (v1.1.21). Zenodo. https://doi.org/10.5281/zenodo.18714417 [84]
  - Levinson, M.A., et al. 2026 - FAIRSCAPE Server (v1.0.32). Zenodo. https://doi.org/10.5281/zenodo.18244417 [85]
- **RO-Crate Validation Classes:**
  - Niestroy, J., et al. 2026 - FAIRSCAPE Pydantic Models (v1.0.1). Zenodo. https://doi.org/10.5281/zenodo.18234523 [72]
  - Leo, S., et al. 2025 - ro-crate-py (0.14.2). Zenodo. https://doi.org/10.5281/zenodo.17342107 [86]
- **LinkML-based Semantic Modeling and Translation:**
  - S.A.T. Moxon, H. Solbrig, N.L. Harris, P. Kalita, M.A. Miller, S. Patil, K. Schaper, C. Bizon, J.H. Caufield, S. Cirujano Cuesta, C. Cox, F. Dekervel, D.M. Dooley, W.D. Duncan, T. Fliss, S. Gehrke, A.S.L. Graefe, H. Hegde, A. Ireland, J.O.B. Jacobsen, M. Krishnamurthy, C. Kroll, D. Linke, R. Ly, N. Matentzoglu, J.A. Overton, J.L. Saunders, D.R. Unni, G. Vaidya, W.-M.A.M. Vierdag, T. Putman, LinkML Community Contributors, O. Ruebel, C.G. Chute, M.H. Brush, M.A. Haendel, C.J. Mungall, LinkML: A Linked Open Data Modeling Language, (2026). https://doi.org/10.5281/ZENODO.5703670. [87]

### Datasets

#### NICU Highly Comparative Time Series Analysis

This data and its provenance (evidence) graph were packaged by an early version of FAIRSCAPE, before RO-Crate Packaging was added to bind provenance and other metadata to the datasets. The datasets and provenance were exported from FAIRSCAPE to the University of Virginia Dataverse instance for long-term archival.

○ Niestroy, J, et al. 2021. Evidence For: Discovery of Signatures of Fatal Neonatal Illness In Vital Signs Using Highly Comparative Time-series Analysis. University of Virginia, 2021-03-31, 2021. https://doi.org/10.18130/V3/HHTAYI [88]
○ Niestroy, J., et al. 2021. *Replication Data For: Discovery of Signatures of Fatal Neonatal Illness In Vital Signs Using Highly Comparative Time-series Analysis*. University of Virginia, 2021-03-26, 2021. https://doi.org/10.18130/V3/VJXODP [89]

- **Bridge2AI Functional Genomics (Cell Maps for Artificial Intelligence - CM4AI) data releases**. The RO-Crate packages were exported from FAIRSCAPE to the University of Virginia Dataverse instance for long-term archival. Inspection of the packages shows the evolution of both data and metadata completeness during the program.
  - Clark T, Schaffer L, Obernier K, et al. Cell Maps for Artificial Intelligence - Data Release V1. University of Virginia Dataverse. https://doi.org/10.18130/V3/DXWOS5 [90]
  - Clark T, Parker J, Al Manir S. Cell Maps for Artificial Intelligence - March 2025 Data Release (Beta) V1. University of Virginia Dataverse. https://doi.org/10.18130/V3/B35XWX [91]
  - Clark T, Parker J, Al Manir S. Cell Maps for Artificial Intelligence - June 2025 Data Release (Beta) V3. University of Virginia Dataverse. https://doi.org/10.18130/V3/F3TD5R [92]
  - Clark T, Parker J, Al Manir S, et al. Cell Maps for Artificial Intelligence - October 2025 Data Release (Beta) V3. University of Virginia Dataverse. https://doi.org/10.18130/V3/K7TGEM [93]

